# An optimized method for the isolation of urinary extracellular vesicles for molecular phenotyping: detection of biomarkers for radiation exposure

**DOI:** 10.1101/2022.01.28.477909

**Authors:** Charles P. Hinzman, Meth Jayatilake, Sunil Bansal, Brian L. Fish, Yaoxiang Li, Yubo Zhang, Shivani Bansal, Michael Girgis, Anton Iliuk, Xiao Xu, Jose A. Fernandez, John H. Griffin, Elizabeth A Ballew, Keith Unger, Marjan Boerma, Meetha Medhora, Amrita K. Cheema

## Abstract

**Background:** Urinary extracellular vesicles (EVs) are a source of biomarkers with broad potential applications across clinical research, including monitoring radiation exposure. A key limitation to their implementation is minimal standardization in EV isolation and analytical analysis methods. Further, most urinary EV isolation protocols necessitate large volumes of sample. This study aimed to compare and optimize isolation and analytical methods for EVs from small volumes of urine.

**Methods:** 3 EV isolation methods were compared: ultracentrifugation, magnetic bead-based, and size-exclusion chromatography from 0.5 mL or 1 mL of rat and human urine. EV yield and mass spectrometry signals (Q-ToF and Triple Quad) were evaluated from each method. Metabolomic profiling was performed on EVs isolated from the urine of rats exposed to ionizing radiation 1-, 14-, 30- or 90-days post-exposure, and human urine from patients receiving thoracic radiotherapy for the treatment of lung cancer pre- and post-treatment.

**Results:** Size-exclusion chromatography is the preferred method for EV isolation from 0.5 mL of urine. Mass spectrometry-based metabolomic analyses of EV cargo identified biochemical changes induced by radiation, including altered nucleotide, folate, and lipid metabolism. We have provided standard operating procedures for implementation of these methods in other laboratories.

**Conclusions:** We demonstrate that EVs can be isolated from small volumes of urine and analytically investigated for their biochemical contents to detect radiation induced metabolomic changes. These findings lay a groundwork to develop future methods to monitor response to radiotherapy and can be extended to an array of molecular phenotyping studies aimed at characterizing EV cargo.

## INTRODUCTION

Extracellular vesicles (EVs) are fast becoming a preferential platform for liquid biopsy-based biomarker discovery owing to rich molecular cargo contents. EVs are released from cells of all tissue types during normal biological processes and have high potential as a source of low abundance biomarkers for a wide array of pathophysiologies (*1, 2*). Given that EVs carry internal cargo, analytical platforms like mass spectrometry are crucial for elucidating their biochemical content. However, studies leveraging techniques such as metabolomics and lipidomics to analyze EV content have so far remained relatively limited.

We have previously examined the role of plasma derived EVs as biomarkers for radiation injury (*3, 4*). Whole-body exposure to acute, large doses (>2 Gy) of ionizing radiation (IR) can potentially be lethal if not diagnosed and treated expeditiously (*5, 6*). Damage to the vascular endothelium may occur as early as 24 hours’ post-exposure, affecting most commonly the gut and bone marrow (*7, 8*). Cumulative IR exposure for the treatment of various cancers also puts patients at risk of normal tissue toxicity (*9–11*). Late effects of IR injury may appear several months delayed, potentially causing life-threatening injuries to the brain, heart, lung, and kidneys. When considering a non-invasive approach for the identification of IR injury biomarkers, we and others have studied plasma, serum, urine, and saliva (*12–14*). However, these matrices may contain significant background, obscuring biomarkers of biological importance.

Urine, as a noninvasive biological matrix, contains a plethora of molecular information indicating rapid changes that occur in various physiological conditions. Urine has been studied as a source of biomarkers related to IR exposure (*15*). Urinary EVs have also shown promise as biomarkers for kidney disease, urological cancers, and even neurologic disease (*16–18*). However, many reported protocols for the isolation of EVs from urine necessitate the use of large volumes of sample (*19, 20*). To our knowledge, no groups have studied the utility of urinary EVs as a source for biomarkers of radiation damage. The goal of this study was to compare, optimize, and validate a method of EV isolation from small volumes of urine, with an emphasis on downstream LC-MS/MS-based metabolomic and lipidomic analyses. We compared three methods of EV isolation: ultracentrifugation (currently one of the most widely used methods with relatively low throughput, generally requiring large volumes (>10 mL) of urine); a magnetic bead-based method (MBB); and a size exclusion chromatography (SEC)-based method. Our benchmarks of success of EV isolation included yield and purity for a given volume, compatibility with LC-MS analyses, and scalability to support large batches of samples. Following extensive characterization to confirm EV enrichment and small molecule profiling, our results showed that SEC-based urinary EV isolation gave optimal yields with 0.5 mL of rat urine and resulted in high quality LC-MS data. The method was validated using a pilot study and later tested using a larger cohort of test samples utilizing the WAG/RijCmcr rat model of leg-out partial body irradiation (leg-out PBI). This sophisticated model is the only one available to display multiple relevant organ sequelae (gastrointestinal, bone marrow, pulmonary, heart, brain, and kidney radiation injuries) observed after a large single radiation exposure. Finally, we demonstrated the clinical utility of our EV analytical approach in a pilot study of human urine samples from patients receiving thoracic radiation therapy (RT).

To our knowledge, this study is the first comparison of EV isolation methods from small volumes of urine, the first report of a comprehensive method for mass spectrometric analysis of the small molecule and lipid content from EVs isolated from small volumes of urine, and the first report evaluating the efficacy of urinary EVs as biomarkers for radiation injury. The availability of reproducible methods of EV isolation from low volumes of urine that are compatible with mass spectrometry methodologies is an unmet need for enabling biomarker discovery, and for studying disease onset, disease progression and patient response to RT. The reported SEC-based small volume urinary EV isolation method can be extended to other types of biomolecular analyses; to this end, we have developed and provided a bulleted optimized standard operating procedure (SOP) to enable easy implementation in other laboratories.

## MATERIALS AND METHODS

### Animal care and irradiation protocols

All animal protocols were approved by the Institutional Animal Care and Use Committees (IACUC) at the Medical College of Wisconsin, Milwaukee. WAG/RijCmcr female rats were irradiated at 11-12 weeks of age (~155 grams). Two groups of rats were randomized for this study: I) No irradiation (n=5); II) 13 Gy leg-out partial body irradiation (leg-out PBI) (n=5). A subset of 4 urine samples from these rats were used for optimizing EV isolation. To test the effect of radiation on EV cargo composition, a separate cohort of rats was randomized into 1) No irradiation, vehicle (n=8), or 2) 13 Gy leg-out PBI (n=10). The protocol used for leg-out PBI was the same as reported in previous studies (*21, 22*) and details can be found in the **Supplementary Materials and Methods**. Urine was collected 24 hours, 14, 30, or 90 days after irradiation. Urine was centrifuged at 13,000 rpm at 4 °C for 10 minutes and the supernatant stored frozen at −80 °C until analyses.

### Human urine collection

The biospecimen collection study was approved by the Georgetown University-MedStar Health Institutional Review Board (IRB 2013-0049) and eligible patients underwent signed informed consent. Eligibility included patients receiving thoracic radiation therapy. Enrolled patients included 12 patients with non-small cell lung cancer and 1 patient with thymoma who were treated 35-66 Gy. A clean catch urine sample was collected prior to radiation therapy, following the last fraction of treatment, and approximately 6 weeks after treatment. For the current study, we used samples from 5 patients pre- and immediately post-RT.

### Urinary EV isolation

We have submitted all relevant data of our experiments to the EV-TRACK knowledgebase (EV-TRACK ID: EV210076) (*23*). EVs were isolated from urine using 3 independent methods (I) ultracentrifugation (UC) with filtration, (II) size exclusion chromatography (SEC) proceeded by filtration/concentration steps, and (III) a proprietary magnetic bead-based isolation method. We compared two different initial volumes of urine (0.5 mL or 1 mL). Detailed protocols and checklists can be found in **Supplementary Files** and **Supplementary Materials and Methods**.

### Nanoparticle Tracking Analyses (NTA)

NTA was performed using a NanoSight NS300 (Malvern Panalytical) equipped with a high sensitivity sCMOS camera, 532 nm laser, and automatic syringe pump. Detailed methods can be found in **Supplementary Materials and Methods**.

### EV immunoblot array

To examine expression of accepted EV specific protein markers in isolated urinary EV samples, we performed the Exo-check Antibody Array (System Biosciences, Palo Alto CA, #EXORAY210A). 30 μg of EV protein was aliquoted and immunoblots were processed and developed according to the manufacturer’s protocol.

### Cryogenic Electron Microscopy

EV samples resuspended in 1 x PBS were sent to the Molecular Electron Microscopy Core at the University of Virginia for analysis. Detailed methods can be found in the **Supplementary Materials and Methods.**

### Mass spectrometry solvents and reagents

All solvents were LC-MS grade. Catalog numbers for all solvents, reagents and standards can be found in the included SOPs in **Supplemental Files**.

### Urinary EV metabolomics using UPLC-QToF-MS

Detailed sample preparation methods and MS data acquisition and processing can be found in **Supplemental Materials and Methods**. **Urinary EV LC-MS/MS polar and lipidomics analysis, data acquisition:** Methods were developed for quantitation using a QTRAP 5500 LC-MS/MS System (Sciex). Detailed methods, protocols and checklists can be found in **Supplemental Files** and **Supplementary Materials and Methods.**

### Statistical analyses and visualization

Prior to statistical analyses, MS data were normalized as described above for each dataset. Binary comparisons were done using paired or unpaired t-tests as appropriate, and fold changes were calculated to identify differences in metabolite levels after radiation. All analyses were performed in R (v 4.0.3). Figures were created using R, GraphPad Prism (v. 9.0) and BioRender (www.BioRender.com).

## RESULTS

### Size exclusion chromatography is the preferred method for isolating EVs from low volumes of urine for mass spectrometry-based metabolomics analysis

To optimize a reliable method for EV isolation from small volumes of urine, we first compared the yield and purity of EVs isolated from two initial volumes of rat urine (0.5 mL or 1 mL) using three independent isolation methods (**Fig. 1**): ultracentrifugation with filtration (UC), size exclusion chromatography (SEC) and a proprietary magnetic bead-based commercial method (MBB). After isolation, we measured the size distribution and concentration of EVs from samples using nanoparticle tracking analysis (NTA). NTA showed all three methods resulted in EVs of similar size (**Suppl. Fig. 1**), though there were differences in the average size of detected particles (**Suppl. Table 1**). We found that the MBB method provided the highest concentration (particles/mL) of EVs at both initial volumes, but SEC resulted in the highest number of total EVs (yield) based upon the entire volume of working solution obtained by each method (**Suppl. Fig. 1**). UC was least efficient (**Suppl. Fig. 1**). Typical yields from 0.5 mL of rat urine using the SEC method ranged from 1.42E9 to 2.82E9 particles/mL, while 1.0 mL of urine yielded between 4.66E9 to 1.13E10 particles/mL (**Suppl. Table 1**). We next used cryogenic electron microscopy (Cryo EM) to confirm EV enrichment and analyze the biophysical properties of isolated vesicles. We found that UC and SEC yielded EVs with of varied sizes, though most within the expected range of < 200 nm in diameter (**Suppl. Fig. 2**). The MBB method, however, seemed to yield smaller vesicles of a more uniform size, an observation other groups have made using polymer-based EV isolation methods (**Suppl. Fig. 2**).

**Figure 1 -.**
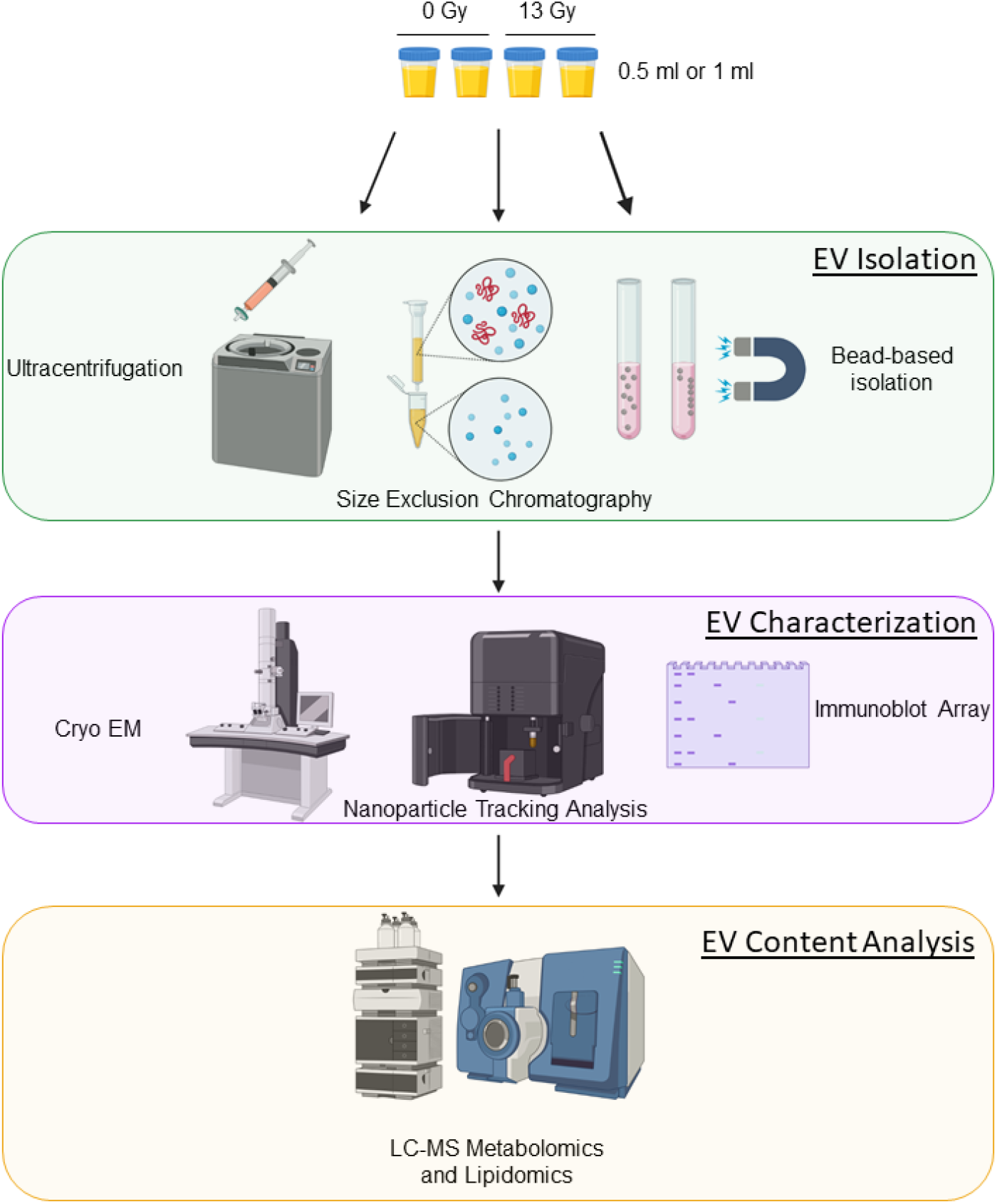
Extracellular vesicle (EV) isolation method comparison workflow. EVs were isolated from 2 volumes of urine (0.5 mL or 1 mL) from rats exposed to either 0 Gy (sham) or 13 Gy, X-irradiation, using 3 independent methods; ultracentrifugation with filtration (UC), size-exclusion chromatography (SEC) or a proprietary magnetic bead-based method (MBB). EV isolates were then characterized using cryogenic electron microscopy, nanoparticle tracking analysis and immunoblot array. Finally, the biochemical content of EVs was evaluated using untargeted quadrupole time of flight (QTOF) mass spectrometry for both overall detection signals and potential contaminants.

To compare resultant mass spectrometry data quality, we next investigated small molecule profiles of EVs isolated via each method separately using quadrupole time of flight mass spectrometry coupled with ultra-performance liquid chromatography (QToF-LCMS)-based untargeted metabolomics analysis. The total number of features (*m/z* and retention time pairs) identified for UC (0.5 mL = 4,837 and 1 mL = 5,141) and SEC (0.5 mL = 5,031 and 1 mL = 5,096) were comparable. EVs isolated using the MBB method had a significantly higher number of detected features (0.5 mL = 7,450 and 1 mL = 7,135) (**Fig. 2A**). Though increasing the starting volume of urine to 1 mL increased EV yield and concentration, the number of detected features only marginally increased (**Fig. 2A**). Interestingly, examination of total ion current chromatograms (TICs) and Manhattan plots revealed a cluster of features which were uniquely detected within the MBB samples (outlined in red, **Fig. 2B-C**). Further investigation of the unique chromatographic peaks in MBB samples revealed *m/z* patterns which were consistent with polymer contaminants. EV preparation using the UC and SEC methods generated MS spectra free of contaminants with profiles similar to each other. We did not observe significant differences in the total mass spectrometry signal between EVs derived from either 0.5 mL or 1 mL of urine (**Fig. 2B-C**). There was significant overlap in the number of unique features identified by each isolation method, with both 0.5 mL and 1 mL of urine (**Figs. 2D-E**). Considering the high EV yield, lack of contaminating material, quality of mass spectrometry data and high throughput capability we determined SEC as the optimal method for isolating EVs from 0.5 mL of urine.

**Figure 2 -.**
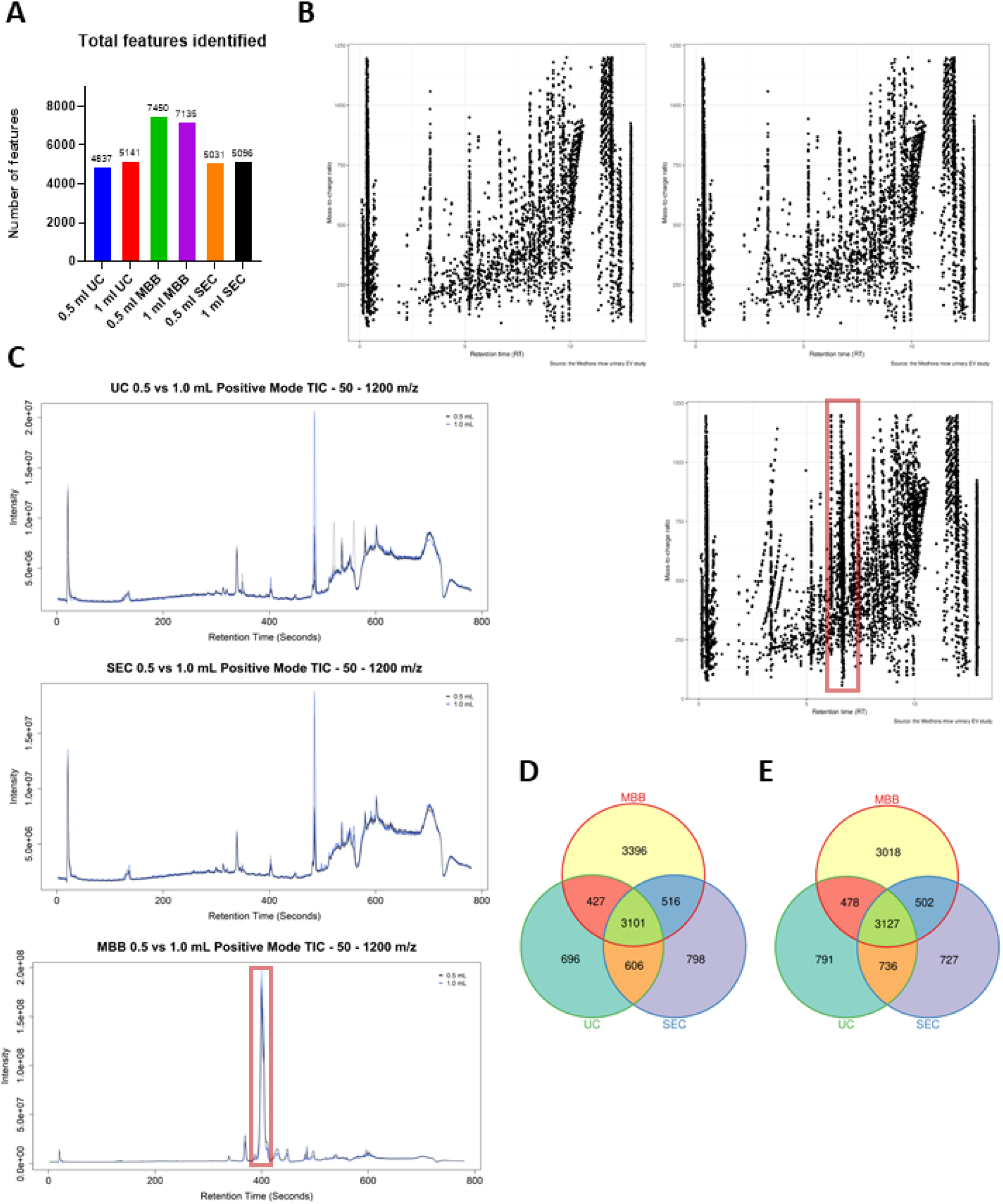
Mass spectrometry analysis of EV isolation methods. **(A)** Total number of features (positive and negative ionization) identified by each isolation method, at each starting volume of urine. **(B)** Manhattan plots showing each feature by mass-to-charge ratio (*m/z*, y-axis) and retention time (*rt*, x-axis). Highlighted area in red shows distinct features in MBB samples which were not detected in EVs isolated by either UC or SEC. **(C)** Total ion chromatogram (TIC) plots from positive ionization mode of EV samples isolated by UC (top), SEC (middle) and MBB (bottom). Area highlighted in red shows distinct signals detected in MBB samples which were not detected in either UC or SEC samples. **(D-E)** Venn diagrams showing the number of unique features detected in EV samples isolated by each isolation method from **(D)** 0.5 mL or **(E)** 1 mL starting volume of urine.

The observation of “classical” EV-markers in antibody array immunoblots was used to ascertain isolation and enrichment of *bona fide* EVs. In accordance with MISEV guidelines (*24*), we performed a detailed characterization of the EV preparations. Firstly, we evaluated the expression of transmembrane proteins (such as CD63 and CD81) as well as cytosolic proteins (such as TSG101, ALIX) in the urinary EV preparations. We found that the urinary EV preparations were positive for known EV markers including cluster of differentiation 63 (CD63), cluster of differentiation 81 (CD81), tumor susceptibility gene 101 (TSG101), ALG-2-interacting Protein X (ALIX), intracellular adhesion molecule (ICAM), Annexin5, epithelial cell adhesion molecule (EpCAM), and flotilin1 (Flot1) (**Suppl. Fig 2**). Importantly, pooled samples of fractions not expected to contain EVs showed no enrichment in these targets (**Suppl. Fig 2**).

### Pilot study validates EV isolation method and potential utility of urinary EVs as a source of small molecule radiation biomarkers

We next performed a pilot study to investigate the utility of urinary EVs as biomarkers of radiation injury. We isolated EVs using our SEC method from 0.5 mL of urine from a small cohort of rats (n = 5 per group) either sham irradiated or exposed to 13 Gy leg-out PBI and performed QToF-LCMS to characterize their small molecule profiles (**Fig. 3A**). Principal component analysis (PCA) demonstrated distinct separation in the small molecule profiles of EVs from irradiated rats (**Fig. 3B**). Visualization using a volcano plot identified many features which were significantly dysregulated (**Fig. 3C**). A heatmap also showed distinct differential expression patterns between irradiated and sham irradiated EV samples (**Fig. 3D**). Overall, we identified a total of 72 features which were significantly altered (FDR adjusted *p*-value < 0.05) in the radiation group (**Suppl. Table 2**). Ultimately, we putatively annotated 21 of these features covering a broad range of endogenous metabolites such as lipids, prostaglandins, peptides or amino acid derivatives, and small molecules such as adrenaline (**Suppl. Table 3**).

**Figure 3 -.**
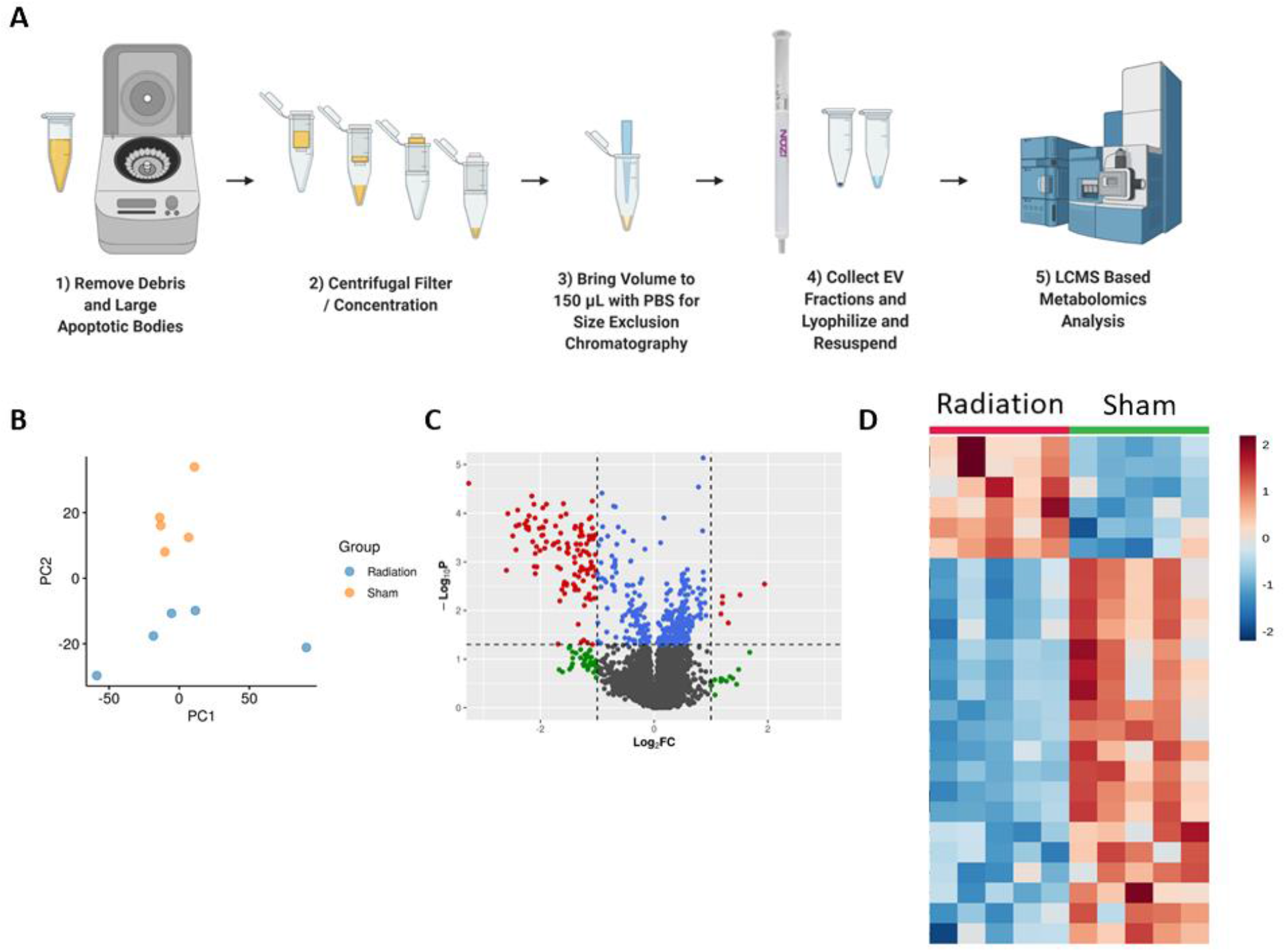
EVs isolated from small volumes of urine demonstrate potential as a source of biomarkers for ionizing radiation exposure. **(A)** Workflow showing EV isolation process from small volumes of urine to untargeted QTOF metabolomics analysis. **(B)** Principal Component Analysis (PCA) plot demonstrates distinct separation between EVs isolated from sham irradiated rats (yellow) vs. rats exposed to 13 Gy irradiation (blue). **(C)** Volcano plot reveals a significant number of detected features are significantly dysregulated. Each dot represents a feature (*m/z* and *rt* pair) detected by QTOF-MS. Grey = no significance, green = significant by fold change (>2 or <0.5), blue = significant by FDR-adjusted *p*-value (<0.05) and red = significant by both FDR adjust *p*-value (<0.05) and fold change (>2 or <0.5). **(D)** Heatmap of features identified in irradiated (left) and sham irradiated (right) urinary EVs demonstrating distinct signatures. Color represents fold change with red indicating upregulation and blue indicating downregulation.

### Urinary EVs allow for the identification of radiation biomarkers in a large rat cohort

This pilot study confirmed the potential of urinary EVs as a source of biomarkers for radiation exposure. To build on these findings, we applied these methods to a larger cohort of 18 rats and a total of 72 rat urine samples to study biochemical profiles of urine derived EVs obtained from rats exposed to 13 Gy leg-out PBI (**Fig. 4A**).

**Figure 4 -.**
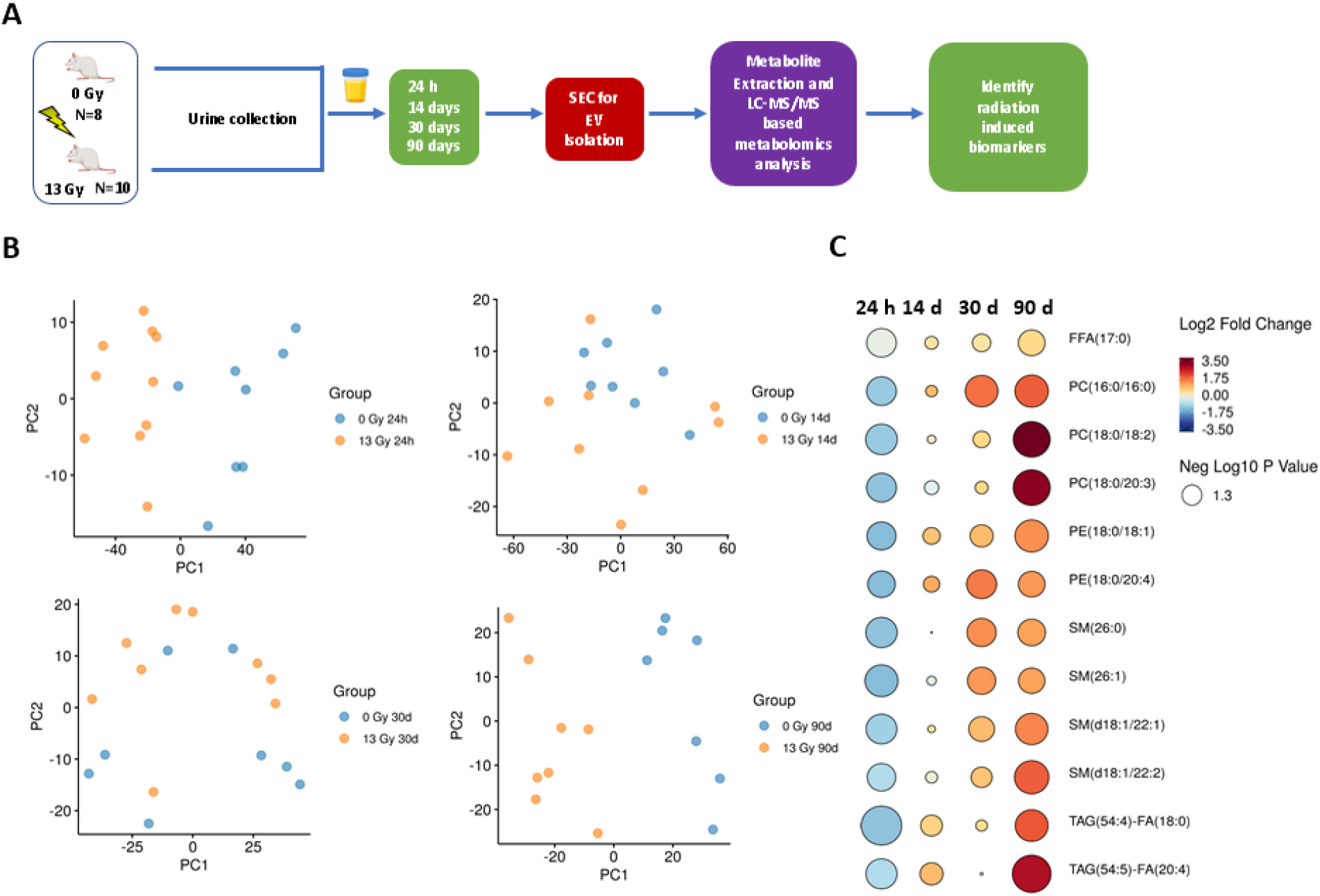
Validation of the potential for EVs isolated from small volumes of urine to serve as biomarkers for ionizing radiation exposure. **(A)** Abbreviated experimental design investigating utility of urinary EVs as a source of radiation biomarkers in rats exposed to 13 Gy ionizing radiation. **(B)** Principal Component Analysis (PCA) plots demonstrate clear separation between EVs isolated from mice exposed to 0 Gy (blue) and 13 Gy (yellow) irradiation 1-, 14-, 30- and 90-days post-irradiation. **(C)** Rain drop plot showing distinct down-regulation of specific lipid species in the acute (24-hour) time frame post-irradiation, followed by recovery and upregulation of these species 14-, 30- and 90-days post-irradiation, compared to control rats.

In accordance with our observations in the pilot study, LC-MS/MS-based targeted metabolomics and lipidomics revealed robust changes in urinary EV profiles following irradiation. Visualization using PCA showed distinct separation between sham and irradiated groups, starkest 24 hours, and 90 days post-irradiation (**Fig. 4B**). No metabolites or lipids were significantly dysregulated (FDR adjusted *p*-value < 0.05) at either 14 or 30-days post-irradiation. Interestingly, the majority of the significantly altered molecules (FDR adjusted *p*-value < 0.05) post-irradiation were lipids (**Suppl. Table 4**). 24 hours post irradiation, several lipid classes were downregulated including triglycerides (TAG), phosphatidylcholines (PC), sphingomyelins (SM), hexosyl ceramides (HCER), free fatty acids (FFAs), lysophosphatidylcholines (LPCs) and phosphatidylethanolamines (PE) (**Fig. 4C, Suppl. Table 4**). By 90 days post-irradiation, many of these lipid species had reversed in their relative abundance with several lipids showing a significant upregulation compared to sham irradiated rats (**Fig. 4C**). The majority of upregulated lipids 90 days post-irradiation were TAGs.

### Urinary EVs demonstrate clinical utility for identifying RT biomarkers in human urine samples

Since the goal of our method comparison is to ultimately develop urinary EV-based biochemical analyses to understand how patients respond to RT in the clinic, we next sought to validate our methods in human urine samples. In a pilot study of 5 thoracic cancer patients receiving RT as a part of their treatment regimen, we collected urine pre- and immediately post-RT, and isolated EVs using the above SEC-based method. First, we validated enrichment of urine EVs using immunoblot and NTA, as described previously (**Suppl. Fig. 3**). We next leveraged our UPLC-QToF-MS platform to analyze the small molecule profiles within our human urine EV samples. Similar to our findings in rat urine EVs, we detected a substantial number of features (4,362 in ESI+ and 3,111 in ESI-) with low noise TICs (**Fig. 5A**). To identify biologically relevant altered metabolites, we used LC-MS/MS targeted metabolomics to quantitate the biochemical profile of human urinary EVs. Using our targeted panel consisting of 360 multiple reaction monitoring (MRM) transitions, covering 270 polar metabolites. We were able to reliably quantify 175 MRMs, corresponding to 152 metabolites, with coefficients of variation less than 0.2, indicating stable and reliable quantification of those metabolites (**Suppl. Table 5**).

**Figure 5 -.**
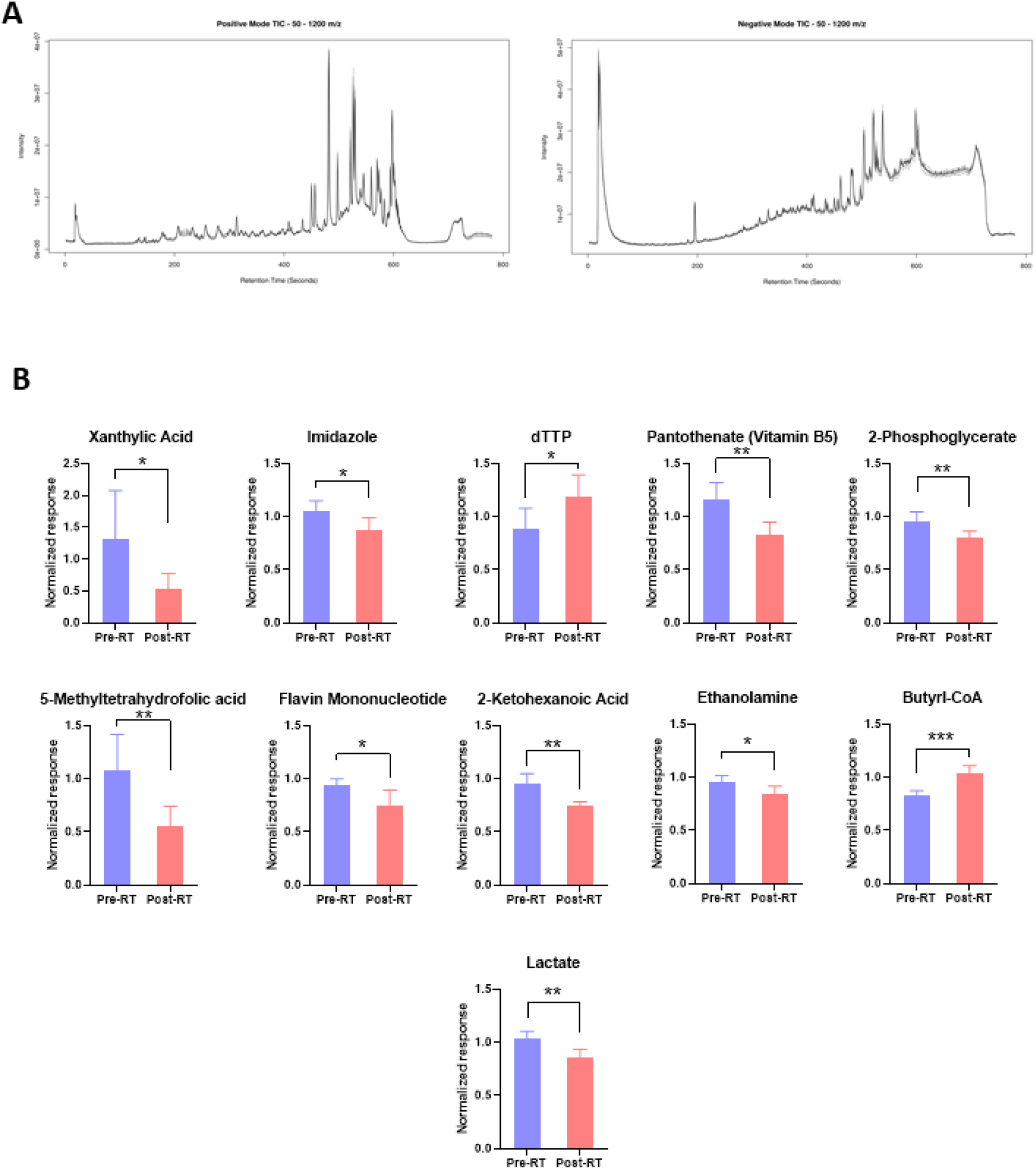
Human urine EVs can be used as a biological matrix to identify the effects of radiation exposure. **(A)** Total ion chromatogram (TIC) plots of human urine EV samples generated by UPLC-QToF-MS in positive (left) and negative (right) ionization mode. **(B)** Metabolites with significant expression changes in human urine EVs post-radiotherapy, quantified using LC-MS/MS. *P*-values: * = < 0.05, ** = < 0.01, *** = < 0.001.

We next sought to identify whether these metabolite profiles significantly changed post-RT. Interestingly, we found 11 metabolites which were stably quantified and significantly altered post-RT in this pilot study (**Table 1**, **Fig. 5B**). Some of these metabolites are involved in nucleotide (Xanthylic acid, imidazole and dTTP) and folate metabolism (5-methyltetrahydrofolic acid and flavin mononucleotide). These changes may implicate signs of DNA damage and/or impaired DNA synthesis upon irradiation.

**Table 1 -.**
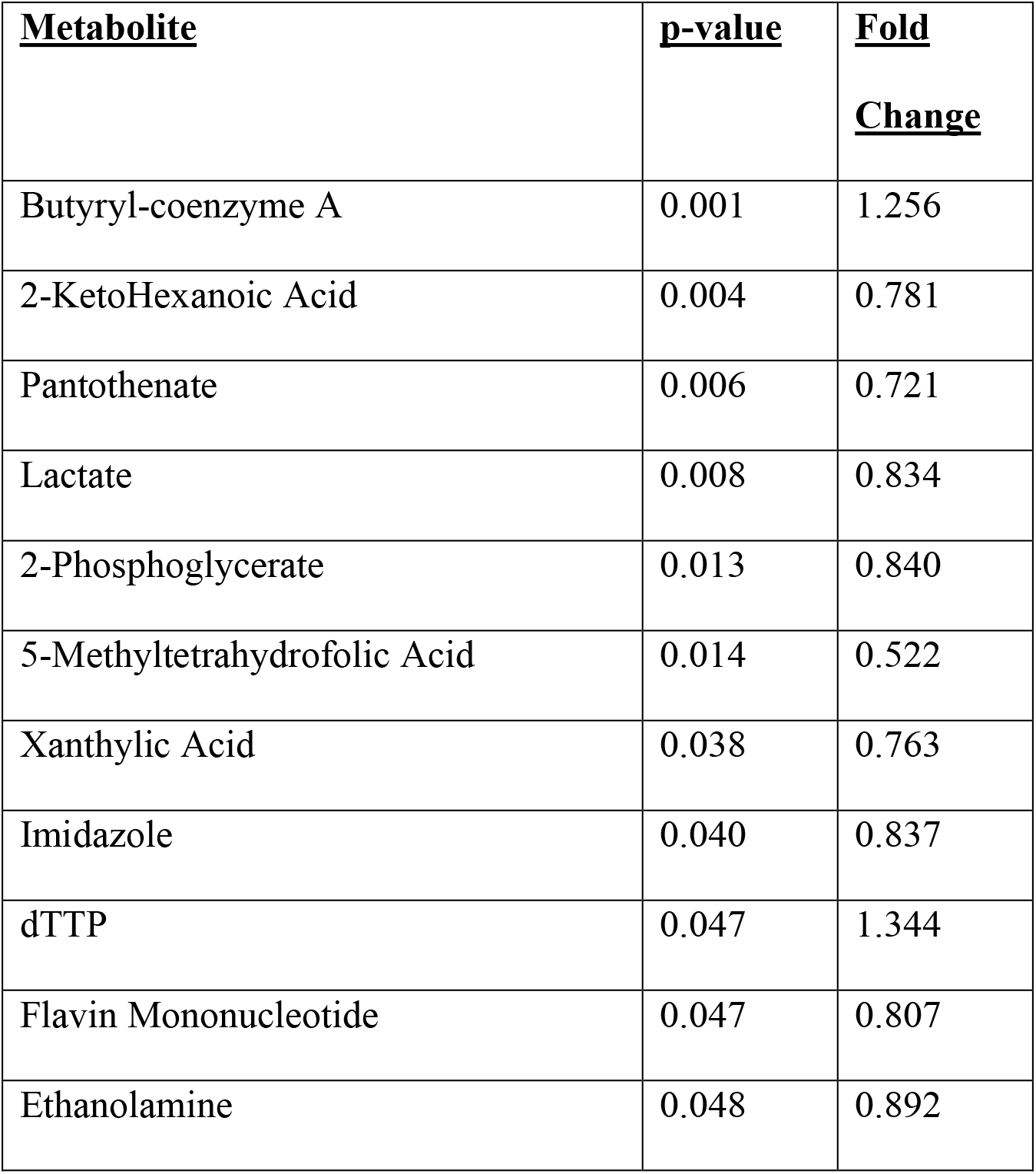
Significantly dysregulated metabolites quantified in human urinary EVs using LC-MS/MS. List of metabolites that were significantly altered (*p*-value < 0.05) in human patient urinary EVs post-RT.

## DISCUSSION

Whole body exposure to high doses of IR can potentially be lethal if radiation injury is not diagnosed and treated expeditiously. Radiation-induced organ damage can manifest within days to years after irradiation. When considering a non-invasive approach for the identification of biomarkers of exposure to IR and of radiation-induced organ damage, we and others have studied molecules in plasma, serum, saliva, and urine. However, these matrices are complex with high abundance molecules like albumins, globulins that can obscure the detection of potential biomarkers of biological importance that have relatively lower abundance. Extracellular vesicles are fast becoming a platform for biomarker discovery in radiation research as well as in other pathologies. However, few studies have investigated the use of a metabolomics approach to analyze EVs derived from urine in the context of IR exposure. Furthermore, the dominant protocols for EV isolation from urine require a large (up to 30 mL) amount of starting volume, which may not be feasible for many studies and clinical translation. The aim of this study was to optimize EV isolation from rat urine and assess radiation-induced alterations in urinary EV metabolic content. Given that EVs have shown tremendous potential as a source for biomarkers for an array of diseases and disorders, our overall goal was to develop a robust and reproducible method of EV isolation from small volumes of urine that is compatible with downstream molecular characterization of EV cargo. Importantly, each of our isolation methods yielded a different number of total detected features. Part of this variance may be explained by the inherent differences in non-EV contaminants co-isolated with each method (*25–27*). This is an important consideration particularly when isolating EVs from other biological matrices which may have high protein concentrations, such as plasma or tissue. The ability to successfully isolate EVs from small volumes of urine can be applied beyond metabolomics to miRNA isolation, proteomics, and functional *in vitro* assays. The method presented is consistent and reliable for isolating, quantifying, and characterizing urinary EVs for research and clinical purposes.

We also demonstrate the utility of urinary EVs as an effective source of biomarkers for detecting IR exposure. Previous reports have studied whole urine, whole plasma, and plasma EVs as sources for biomarkers of IR exposure and IR damage (*3, 11, 28–31*). Dyslipidemia has previously been observed after radiation exposure (*3, 12, 32–35*). Given the importance of lipid composition to EV function, this remains an active area of research. We have previously found that post-IR exposure, plasma EVs are enriched in lipid species indicative of an inflammatory response, namely TAGs (*4*). In that previous study, we found that upregulation of TAGs dissipated by 14 days post-irradiation, a finding similar to our results reported here. Xu et. al previously demonstrated that TAGs are upregulated in the plasma at late time points (15-20 weeks) post irradiation (*13*). TAGs were the bulk of lipids dysregulated 90 days post-irradiation, all of which were upregulated. The functional consequences of these changes remain an active area of research for our group.

These methods are also useful for studying human clinical samples. Though we have only performed a pilot study on 10 patients receiving RT as a part of their thoracic cancer treatment regimens, we were able to detect significant metabolite profiles of urinary EVs from extremely small volumes of urine. Importantly, these differences may also be biologically relevant in the context of IR exposure. The same data can also be extrapolated for correlation of EV-metabolites with tumor response to radiation therapy, once follow-up data become available. As reported here, we found significantly altered levels of key metabolites driving folate and nucleotide metabolism. Previous studies have demonstrated functional consequences of folate processing enzymes and altered folate pools imparted by radiation stress (*36*). Activation of DNA damage and repair mechanisms, as well as altered nucleotide metabolism are canonical biological events post-IR exposure, events that may be linked to radio responsiveness depending on tissue type and disease setting (*37–39*). Considering previous reports, with our findings, urinary EVs present themselves as a potentially important biological matrix for monitoring metabolite changes, and ultimately patient response, to RT.

In conclusion, the urinary EV metabolome and lipidome may be a sensitive and specific early indicator of radiation injury and a platform for monitoring patient response to RT. Furthermore, the approaches developed and validated in this study can be easily applied in diverse areas of biomedical research that seek to leverage molecular analyses of urinary EV profiles for gaining insights into disease onset and progression. Finally, of the methods tested herein, SEC was found to be the preferred method for isolating EVs from small volumes of urine for broad-based mass spectrometric analysis.

## Supporting information

Supplemental Methods, Figures, Tables and SOPs

## FUNDING AND ACKNOWLEDGMENTS

This study was supported by three grants and a supplement from NIAID/NIH [U01AI133561, U01AI148308 to AKC and MB and P20 GM109005 to MB] and two grants and a supplement to MM [U01AI133594 and U01AI107305]. Research reported in this publication was also supported by the National Center for Advancing Translational Sciences of the National Institutes of Health under Award Number TL1TR001431. The content is solely the responsibility of the authors and does not necessarily represent the official views of the National Institutes of Health. The authors wish to thank Jayashree Narayanan for coordinating the animal study and Tracy Gasperetti and Dana Scholler for animal care and treatments. The authors would like to acknowledge the Metabolomics Shared Resource as well as the Flow Cytometry & Cell Sorting Shared Resource at Georgetown University which are partially supported by NIH/NCI/CCSG grant P30-CA051008. The authors would also like to acknowledge Dr. Kelly Dryden at the Molecular Electron Microscopy Core at the University of Virginia.

## DECLARATION OF INTEREST STATEMENT

The authors confirm that there are no known conflicts of interest associated with this publication.

## DATA AVAILABILITY STATEMENT

All datasets generated in the publication of this study are available by reasonable request to the corresponding author.

## Notes

### Competing Interest Statement

The authors have declared no competing interest.

