## Supplemental Methods, Figures, Tables and SOPs for "An optimized method for the isolation of urinary extracellular vesicles for molecular phenotyping: detection of biomarkers for radiation exposure": bioRxiv-An optimized method for isolation of urinary EVs_SOPs.docx

**Detailed protocols and standard operating procedures for:**

**Updated November 2021**

**Protocol for Isolation of Extracellular Vesicles from Urine using Size Exclusion Chromatography**

A. **Preparation of Urine**

**Reagents and Equipment:**

0.5 mL Urine Samples	2 mL Tube Centrifuge

2 mL Tubes	1 mL Pipette and Tips

If urine is frozen, you will also need:

37 °C Water Bath

**Procedure:**

1. If urine is frozen, thaw in 37 °C water bath.
2. Centrifuge urine at 2,500 x g for 10 minutes to remove debris.
3. Transfer the supernatant to a new 2 mL tube.
B. **Filtration and Concentration of Urine**

**Reagents and Equipment:**

Prepared Urine from 0.5 mL Sample	2 mL Tube Centrifuge

1 mL and 200 µL Pipettes and Tips	1X PBS (14-190-144)

Leur-Slip Syringe (S7510-10)	Millipore Millex 0.1 µm Filter Unit (SLVVR33RS)

100 kDa Millipore Amicon Ultra 0.5	15 mL Conical Tube

Centrifugal Filter Unit (UFC5100)

**Procedure:**

4. Prepare 150 µL of 0.1 µm filtered 1X PBS per sample in a 15 mL tube.
5. Pipette the prepared urine into the 100 kDa Amicon Ultra filter device.
6. Place filter device into filtrate collection tube.
7. Centrifuge to concentrate and filter at 14,000 x g for 5 minutes at room temperature.
8. Remove filter from filtrate collection tube.
9. Place filter upside down in a new collection tube then centrifuge at 1000 x g for 2 minutes at room temperature.
10. Remove the empty filter then bring up volume in the collection tube to 150 µL with 1X PBS.
C. **Isolation of Urinary EVs Using Size Exclusion Chromatography**

**Reagents and Equipment:**

150 µL Concentrated and Filtered Urine Samples	1 mL and 200 µL Pipettes and Tips

2 mL Tubes	1X PBS (14-190-144)

Leur-Slip Syringe (S7510-10)

Izon qEV Single 70 nm Size Exclusion	Izon Automatic Fraction Collector or Support

Chromatography Columns (SP2)	Stand and Clamp

**Procedure:**

11. Set up Izon qEV single size exclusion chromatography column on fraction collector or stand
12. Remove the cap at the bottom of the column then remove the cap on the top.
13. As the column drips, add a total of 4 mL (4 column volumes) of 0.1 micron filtered 1X PBS until the column stops dripping.
14. Load the 150 µL sample onto the column followed by 0.1 micron filtered 1X PBS. The column will begin dripping immediately with the void volume (total of 1 mL).
15. After 1 mL has been collected, the next 600 µL of eluent will be the EV fraction.
16. Continue adding PBS until ass desired fractions are collected.

**Protocol for Analyte Quantitation of Urinary Extracellular Vesicles Using Sciex QTRAP 5500**

A. **Preparation of Urinary EVs for LC-MS/MS Polar Data Acquisition**

**Reagents and Equipment:**

SEC Urine EV Samples	2 mL Tubes

Dry Ice	37 °C Water Bath

Sonicator	Vortex Mixer

200 µL Pipette and Tips	LC-MS Grade Water (W6-4)

LC-MS Grade Methanol (A456-4)	LC-MS Grade Isopropanol (A461-4)

Debrisoquine (IS) (D1306100MG)	4-Nitrobenzoic Acid (IS) (146657250MG)

4 °C Refrigerator	-20 °C Freezer

2 mL Tube Centrifuge	LC-MS Vials

**Procedure:**

1. Aliquot 10 µL of the prepared SEC urine EV samples into a 2 mL tubes.
2. Heat shock the sample by performing 3 repetitions of 30 seconds in dry ice followed by 90 seconds in a 37 °C water bath.
3. Sonicate the samples for 30 seconds.
4. Add 40 µL of a mixture of 35% water, 25% methanol, and 40% isopropanol to each of the samples then vortex.
5. Add 100 µL of a mixture of 50% water and 50% methanol containing 500 ng/mL of each of debrisoquine (in water) and 4-nitrobenzoic acid (in methanol) to be used internal standards in positive and negative modes, respectively.
6. Vortex the samples for 30 seconds then keep at 4 °C for 20 minutes followed by incubation at -20 °C for 20 minutes.
7. Centrifuge the samples at 15,493 x g for 20 minutes at 4 °C then transfer the supernatant to LC-MS vials for LC-MS/MS analysis.
B. **LC-MS/MS Polar Analysis Using Sciex QTRAP 5500**

**Notes:**

This analysis is used to measure 270 endogenous small molecules using a QTRAP 5500 LC-MS/MS.

**Reagents and Equipment:**

Sciex QTRAP 5500	Shimadzu SIL-30 AC Autosampler

Shimadzu LC-30AD Solvent Delivery Unit	CBM-20A Communication Bus

Prepared LC-MS Urine EV Sample	LC-MS Grade Water (W6-4)

LC-MS Grade Acetonitrile (A955-4)	LC-MS Grade Formic Acid (A117-50)

Phenomenex Kinetex 2.6 μm, 100 Å,

100 × 2.1 mm (00A-4723-AN)

**Procedure:**

- 5 µL of the prepared sample is injected onto the Kinetex 2.6 μm 100 Å 100 × 2.1 mm column kept at 30 °C using the Shimadzu SIL-30 AC auto sampler connected to the Shimadzu LC-30AD solvent delivery unit communicating with the Sciex QTRAP 5500 operating in positive and negative ionization modes via the Shimadzu CBM-20A communication bus.
- The binary gradient consists of water with 0.1% formic acid (solvent A) and acetonitrile with 0.1% formic acid (solvent B) at a flow rate of 0.2 mL/min. The gradient is set as follows:
  a. Initial: 100% A, 0% B
  b. 2.1 minutes: 100% A, 0% B
  c. 14.0 minutes: 5% A, 95% B
  d. 15.0 minutes: 5% A, 95% B
  e. 15.1 minutes: 100% A, 0% B
  f. 23.0 minutes – 100% A, 0% B
- The source and gas settings are as follows:

a. Curtain gas: 35 psi
b. Collisionally Activated Dissociation Gas: Medium
c. Ion Spray Voltage: 2500 V Positive, -4500 V Negative
d. Temperature: 400 °C
e. Nebulizing Gas: 60 psi
f. Heater Gas: 70 psi
C. **Preparation of Urinary EVs for LC-MS/MS Lipidomics Data Acquisition**

**Notes:**

For every 100 mL of the 100% isopropanol internal standards are added in the following amounts: 1 mL of EquiSPLASH® LIPIDOMIX®, 250 µL of 1 mg/mL 18:1 Chol (D7) ester, 500 µL of 100 µg/mL 15:0-18:1-d7-PA, 1.2 mL of Free Fatty Acids (FFA) Internal Standard Kit for Lipidyzer™ Platform (5040166)-1 vial out of 5 vials, 1.2 mL Dihydroceramide (DCER) Internal Standard Kit for Lipidyzer™ Platform (5040397)-1 vial out of 5 vials, 1.2 mL of Hexosylceramide (HCER) Internal Standard Kit for Lipidyzer™ Platform (5040398)-1 vial out of 5 vials, 1.2 mL Lactosylceramide (LCER) Internal Standard Kit for Lipidyzer™ Platform (5040399)-1 vial out of 5 vials.

**Reagents and Equipment:**

SEC Urine EV Samples	2 mL Tubes

Dry Ice	37 °C Water Bath

Sonicator	Vortex Mixer

4 °C Refrigerator	-20 °C Freezer

200 µL Pipette and Tips	LC-MS Grade Water (W6-4)

LC-MS Grade Methanol (A456-4)	LC-MS Grade Isopropanol (A461-4)

EquiSPLASH® LIPIDOMIX® (IS) (330731)	18:1 Chol (D7) ester (IS) (700185)

15:0-18:1-d7-PA (IS) (791642C)	2 mL Tube Centrifuge

Free Fatty Acids (FFA) Internal Standard Kit for	Dihydroceramide (DCER) Internal Standard Kit for

Lipidyzer™ Platform (5040166)	Lipidyzer™ Platform (5040397)

Hexosylceramide (HCER) Internal Standard Kit	Lactosylceramide (LCER) Internal Standard Kit

for Lipidyzer™ Platform (5040398)	for Lipidyzer™ Platform (5040399)

LC-MS Vials

**Procedure:**

8. Aliquot 10 µL of the prepared SEC urine EV samples into a 2 mL tubes.
9. Heat shock the sample by performing 3 repetitions of 30 seconds in dry ice followed by 90 seconds in a 37 °C water bath.
10. Sonicate the samples for 30 seconds.
11. Add 40 µL of a mixture of 35% water, 25% methanol, and 40% isopropanol to each of the samples then vortex.
12. Add 100 µL of 100% isopropanol containing the internal standards (see note above).
13. Vortex the samples for 1 minute then keep at 4 °C for 30 minutes
14. Vortex then incubate samples at -20 °C for 2 hours.
15. Centrifuge the samples at 15,493 x g for 20 minutes at 4 °C then transfer the supernatant to LC-MS vials for LC-MS/MS analysis.
D. **LC-MS/MS Lipidomics Analysis Using Sciex QTRAP 5500**

**Notes:**

This analysis is used to measure 21 classes of lipid molecules using a QTRAP 5500 LC-MS/MS. The 21 lipid classes observed are diacylglycerols (DAGs), cholesterol esters (CEs), sphingomyelins (SMs), phosphatidylcholines (PCs), triacylglycerols (TAGs), monoacylglycerols (MAGs), free fatty acids (FFAs), ceramides (CERs), dihydroceramides (DCERs), hexosylceramides (HCERs), lactosylceramides (LCERs), phosphatidylethanolamines (PE), lysophosphatidylcholines (LPCs), lysophosphatidylethanolamines (LPEs), phosphatidic acids (PAs), lysophosphatidic acids (LPAs), phosphatidylinositols (PI), lysophosphotidylinositols (LPIs), phosphatidylglycerols (PGs), acylcarnitines and phosphatidylserines (PSs).

**Reagents and Equipment:**

Sciex QTRAP 5500	Shimadzu SIL-30 AC Autosampler

Shimadzu LC-30AD Solvent Delivery Unit	CBM-20A Communication Bus

Prepared LC-MS Urine EV Sample	LC-MS Grade Water (W6-4)

LC-MS Grade Acetonitrile (A955-4)	LC-MS Grade Formic Acid (A117-50)

Waters XBridge BEH Amide 3.5 μm, 130 Å,

100 × 2.1 mm (186004868)

**Procedure:**

- 5 µL of the prepared sample is injected onto the Waters XBridge BEH Amide 3.5 μm, 130 Å, 100 × 2.1 mm column kept at 35 °C using the Shimadzu SIL-30 AC auto sampler connected to the Shimadzu LC-30AD solvent delivery unit communicating with the Sciex QTRAP 5500 operating in positive and negative ionization modes via the Shimadzu CBM-20A communication bus.
- The binary gradient consists of 5% water and 95% acetonitrile with 10 mM ammonium acetate (solvent A) and 50% water and 50% acetonitrile with 10 mM ammonium acetate (solvent B) at a flow rate of 0.7 mL/min. The gradient is set as follows:
  a. Initial: 100% A, 0% B
  b. 3.0 minutes: 99.9% A, 0.1% B
  c. 6.0 minutes: 94% A, 6% B
  d. 10.0 minutes: 75% A, 25% B
  e. 11.0 minutes: 2% A, 98% B
  f. 13.0 minutes: 0% A, 100% B
  g. 18.6 minutes: 0% A, 100% B
  h. 18.7 minutes: 100% A, 0% B
  i. 24.0 minutes: 100% A, 0% B (end)
- The source and gas settings are as follows:

g. Curtain gas: 30 psi
h. Collisionally Activated Dissociation Gas: Medium
i. Ion Spray Voltage: 5500 V Positive, -4500 V Negative
j. Temperature: 550 °C
k. Nebulizing Gas: 50 psi
l. Heater Gas: 60 psi
