## Supplemental Methods, Figures, Tables and SOPs for "An optimized method for the isolation of urinary extracellular vesicles for molecular phenotyping: detection of biomarkers for radiation exposure": bioRxiv-An optimized method for isolation of urinary EVs_Supplemental Figures and Materials and Methods.docx

**Supplemental Figure 1 –** **(A-C)** Nanoparticle tracking analysis (NTA) data from samples isolated using either **(A)** Ultracentrifugation, **(B)** Size-exclusion chromatography, or **(C)** a magnetic bead-based method. **(D)** Combined bar graph of EV concentration (particles/mL) isolated from each sample using each method. **(E)** Calculated total number of EVs isolated by each method. Total particles arrived at by multiplying raw NTA concentration reading by dilution factor (yielding undiluted particles/mL), then adjusting for final total recovered sample volume.

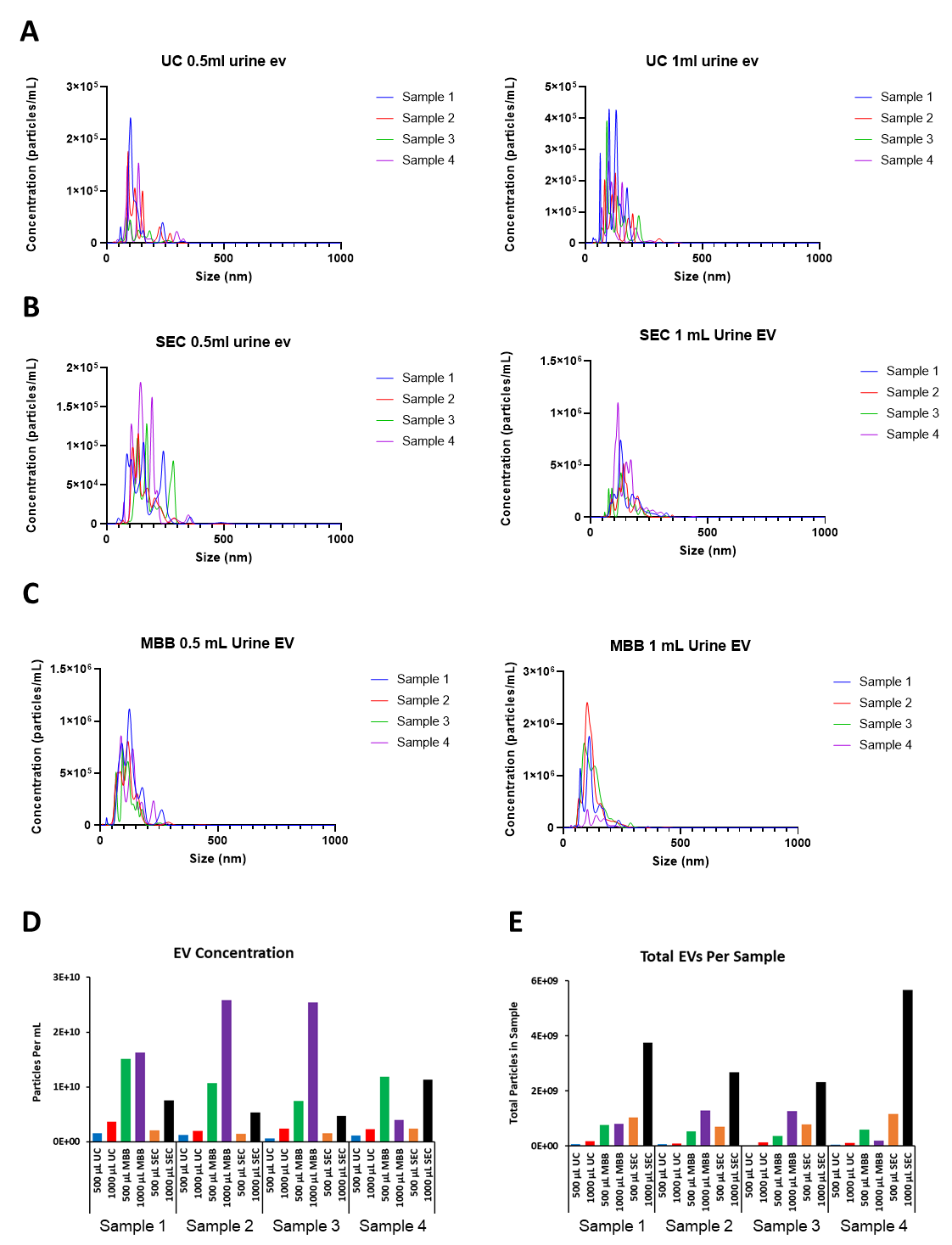

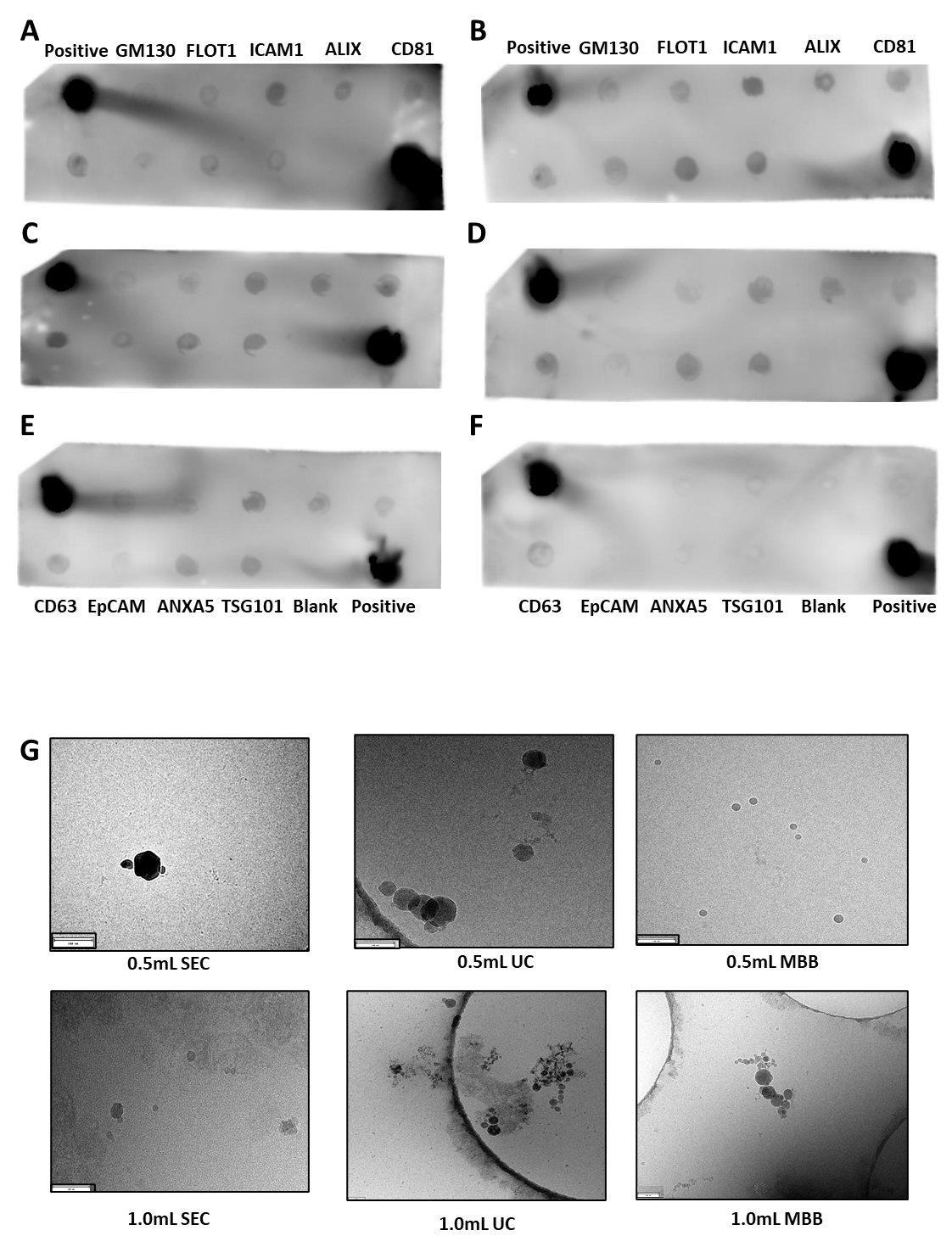
**Supplemental Figure 2 – (A-F)** Immunoblot arrays validation expression of known EV markers from urinary EV samples isolated using SEC and a non-EV fraction after filtration (**F**). **(G)** Cryo-EM photos depicting biophysical characteristics of EVs isolated by each method.

**Supplemental Figure 3 – (A)** Immunoblot array measuring expression of known EV markers in urinary EV samples (left) and non-EV fraction after filtration (right) from a representative human patient urine EV sample. **(B)** Representative nanoparticle tracking analysis (NTA) of human urinary EV samples (N = 5) isolated from patients pre- and post-RT.

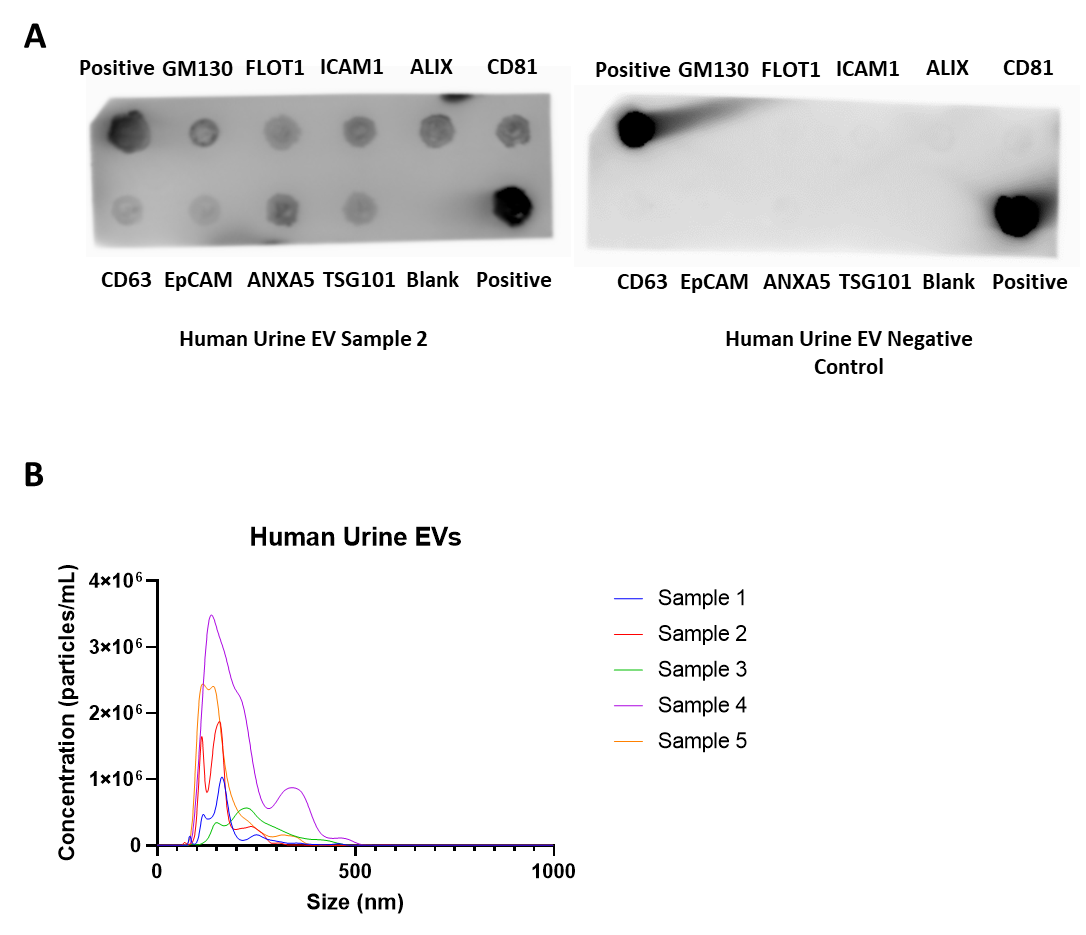

**Supplemental Materials and Methods:**

**Rat irradiation protocol:**

All animal protocols were approved by the Institutional Animal Care and Use Committees (IACUC) at the Medical College of Wisconsin, Milwaukee.

*Animals and groups*: WAG/RijCmcr female rats were irradiated at 11-12 weeks of age (~155 grams). Two groups of rats were randomized for this study: I) No irradiation (n=5); II) 13 Gy leg-out partial body irradiation (leg-out PBI) (n=5). A subset of 4 urine samples from these rats were used for optimizing EV isolation. To test the effect of radiation on EV cargo composition, a separate cohort of rats was randomized into 1) No irradiation, vehicle (n=8), or 2) 13 Gy leg–out PBI (n=10).

Rats were restrained and irradiated without the use of anesthetics. One hind limb of each rat was carefully externalized and shielded with a 0.25-inch lead block. An X-RAD 320KV orthovoltage X-ray system (Precision X-Ray, North Branford, Connecticut) was operated at 320 kVp and 13 mA with a half value layer of 1.4 mm Cu with a dose-rate of 1.69 Gy/min for a total dose of 13 Gy. Radiation was delivered posterior to anterior to the rat. All rats received supportive care including hydration by daily subcutaneous injection of saline 40 mL/kg/day from days 2-10, and antibiotics, and enrofloxacin (10 mg/kg/day) from days 2-14 given in the drinking water. Powdered food was added to the cages from days 35 to 70 after irradiation due to tooth loss that resolved by day 70.

**Urinary EV isolation methods:**

EVs were isolated from urine using 3 independent methods (I) ultracentrifugation (UC) with filtration, (II) size exclusion chromatography (SEC) proceeded by filtration/concentration steps, and (III) a proprietary magnetic bead-based isolation method. We compared two different initial volumes of urine (0.5 mL or 1 mL). All urine samples were thawed at 37 °C before further processing.

**Ultracentrifugation (UC):** Urine was diluted into 1x PBS to a final volume of 30 mL in 50 mL conical tubes. Samples were centrifuged at 1,600 x g for 20 minutes at 25 °C. The supernatants were transferred to UC tubes (#326823, Beckman Coulter) by inversion and samples were centrifuged at 10,000 x g for 20 minutes at 25 °C. Next, supernatants were filtered through 0.22 μm syringe filters (#SLGP033RB, Millipore Sigma) into new UC tubes and centrifuged at 120,000 x g for 70 minutes at 25 °C. Supernatant was discarded by inversion and EV pellets were resuspended in 50 μL of 1x PBS and stored at -80 °C until further use.

**Size-Exclusion Chromatography (SEC):** Urine was aliquoted into new microcentrifuge tubes and centrifuged at 2,500 x g for 10 minutes at 25 °C. Urine was collected and filtered/concentrated through a 100 KDa filter (#UFC5100, Millipore Sigma) by centrifuging at 14,000 x g for 5 minutes at 25 °C. Concentrated urine was collected by inverting the filter into a clean microcentrifuge tube and centrifuging at 1,000 x g for 2 minutes. The recovered volume of urine samples varied between 25-60 μL. Prior to loading samples on SEC columns, sample volumes were adjusted to a final volume of 150 μL using 0.1 μm filtered 1x PBS. Samples were fractionated using 70 nM qEV single columns (SP2, Izon Science) using an automated fraction collector in 200 μL fractions (Izon Science). Fractions containing EVs (#1-#3) were combined, frozen at -80 °C and lyophilized. Samples were then resuspended in 50 μL 1x PBS and stored at -80 °C until further use.

**Magnetic bead-based EV Isolation (MBB):** EVs were isolated according to manufacturer’s protocol. In brief, samples were centrifuged at 2,500 x g for 10 minutes at 25 °C. Supernatant was transferred to fresh tubes, and the magnetic beads supplied were added, at a ratio of 20 μL beads per mL urine. Samples were incubated for 1 hour by end-over-end rotation. Beads were washed 3 times with 0.1% Tween-20 in PBS using a magnetic separator. EVs were eluted using 100 mM triethylamine. Samples were then lyophilized, resuspended in 50 μL 1x PBS and stored at -80 °C until further use.

**Urinary EV metabolomics using UPLC-QToF-MS:** EV samples (50 µL resuspended in 1x PBS) stored at -80 °C were removed from storage. Sample tubes were placed on dry ice for 30 seconds and heat shocked by plunging into a 37 °C water bath for 90 seconds. This cycle was repeated two more times. Samples were then sonicated for 1 minute and incubated on ice for 20 minutes. Next, 75 μL of chilled extraction buffer (35% water, 25% methanol and 40% isopropyl alcohol) containing internal standards (debrisoquine and 4-nitrobenzoic acid) was added. Samples were then vortexed and kept on ice for 20 minutes. Next, 75 μL of chilled acetonitrile was added and samples were vortexed again for 30 seconds and incubated at -20 °C for 20 minutes. Finally, samples were centrifuged at 13,000 x g for 20 minutes at 4 °C and the supernatants were transferred to MS vials for data acquisition.

**UPLC-QToF-MS of Urinary EVs:**

Untargeted analysis of EV samples was accomplished using ultra-performance liquid chromatography (UPLC) on an Acquity UPLC (Waters Corporation) coupled to a Xevo G2-S (Waters Corporation) quadrupole time of flight mass spectrometer (QToF-MS) operating in either positive or negative electrospray ionization. Each sample (1 µL) was injected onto a Waters Acquity BEH C18 130Å, 1.7 µm, 2.1 × 50 mm column set at 40 °C. The gradient mobile phases consisted of 100% water with 0.1% formic acid (solvent A), 100% acetonitrile with 0.1% formic acid (solvent B), 100% isopropanol with 0.1% formic acid (solvent C). Each sample injection was run for 13 minutes at a flow rate for 400 µL/min. The LC gradient conditions with a ramp curve of 6 at each step were as follows: Initial – 95% A, 5% B; 0.5 minutes – 95% A, 5% B; 8.0 minutes – 2% A, 98% B; 9.0 minutes – 11.8% B, 88.2% C; 10.5 minutes - 11.8% B, 88.2% C, 11.5 minutes – 50% A, 50% B; 12.5 minutes – 95% A, 5% B; 13 minutes – 95% A, 5% B.

The column eluent was introduced directly into the G2-S mass spectrometer by electrospray. Positive mode had a capillary voltage of 3.00 kV and a sampling cone voltage of 30 V. Negative mode had a capillary voltage of 2.00 kV and a sampling cone voltage of 30 V. The desolvation gas flow was set to 1000 L/hour and the desolvation temperature was set to 500 °C. The cone gas flow was 25 L/hour and the source temperature was set to 120 °C. The data was acquired in the sensitivity MS mode with a scan time of 0.300 seconds and an interscan time of 0.014 seconds. Accurate mass was maintained by infusing Leucine Enkephalin (556.2771 m/z) in 50% aqueous acetonitrile (0.5 ng/mL) at a rate of 20 µL/min via the Lockspray interface, every 10 seconds. Data was acquired in Centroid mode with a 50.0 to 1200.0 m/z mass range for TOF-MS scanning. Before and after samples run, a mixture of six standards (acetaminophen: m/z 152.0712 [M+H]^+^ / 150.0555 [M−H]^−^, sulfaguanidine: m/z 215.0603 [M+ H]^+^ / 213.0446 [M−H]^−^, sulfadimethoxine: m/z 311.0814 [M+H]^+^ / 309.0658 [M−H]^−^, Val-Tyr-Val: m/z 380.2185 [M+H]^+^ / 378.2029 [M−H]^−^, terfenadine: m/z 472.3216 [M+H]^+^ and leucine-enkephalin: m/z 556.2771 [M+H]^+^ / 554.2615 [M−H]^−^) were run to ensure mass accuracy during data acquisition A number of measures were used to ensure high quality and reproducibility of LC-MS data. The column was conditioned using pooled QC samples, which were injected periodically through the batch to monitor mass accuracy, shifts in retention time and signal intensities as measures of reproducibility.

**UPLC-QToF-MS data pre-processing:** The raw data files obtained from the acquisition on the mass spectrometer were converted into NetCDF files for pre-processing using Waters MassLynx Databridge Software (Waters Corporation). XCMS (*35-37*) was used for data pre-processing while the Isotopologue Parameter Optimization (IPO) package (*38*) was used for XCMS parameter optimization. Normalization was performed with internal standards in both positive and negative mode data. R package was used to perform multivariate analysis. Statistically significant m/z’s with a *p-*value ≤ 0.05 were run on the UPLC-QToF instrument in the MS/MS mode. The MS/MS raw data files were converted to NetCDF files using Waters MassLynx Databridge Software, then further converted to MSP file format using an in-house R package. The m/z’s were thereafter putatively identified with database search by matching MS/MS spectra to entries in the NIST 2017 MS/MS database and by applying the online version of CEU Mass Mediator (CMM) which integrated from METLIN, Human Metabolome Database (HMDB) and LIPID MAPS with a ppm error of less than 10.

**EV LC-MS/MS polar and lipidomics analyses**:

Methods were developed for quantitation using a QTRAP 5500 LC-MS/MS System (Sciex). The polar method was developed to quantitate 270 endogenous small molecules. The lipidomics method was designed to measure 21 classes of lipid molecules, including diacylglycerols (DAGs), cholesterol esters (CEs), sphingomyelins (SMs), phosphatidylcholines (PCs), triacylglycerols (TAGs), free fatty acids (FFAs), ceramides (CEs), dihydroceramides (DCERs), hexosylceramides (HCERs), lactosylceramides (LCERs), phosphatidylethanolamines (PEs), lysophosphatidylcholines (LPCs), lysophosphatidylethanolamines (LPEs), phosphatidic acids (PAs), lysophosphatidic acids (LPAs), phosphatidylinositols (PIs), lysophosphotidylinositols (LPIs), phosphatidylglycerols (PGs) and phosphatidylserines (PSs). Full protocols can be found in Supplementary Files.

**LC-MS/MS data processing:** The data were normalized to respective internal standards and processed using MultiQuant 3.0.3 (Sciex). The quality and reproducibility of LC-MS data was ensured using a number of measures. The column was conditioned using pooled QC samples which were injected every 10 sample injections throughout the sample batch. We also ran NIST plasma samples, prepared using the same sample preparation method as the actual samples, after every 20 samples to monitor instrument variance. Solvent blanks were injected between sets of samples to monitor and ensure there was no sample-to-sample carry-over. The data were pre-processed using a signal/noise ratio >20:1 and retention time (RT) tolerance of 5 seconds. Data quality monitoring-based on coefficient of variation of each feature in pooled QC samples run periodically.

**Cryogenic Electron Microscopy:** EVs resuspended in 1 x PBS (~3.5 μL) were applied to a glow-discharged, perforated 2/1-3C C-Flat carbon-coated grids (Protochips, Raleigh, NC). Samples were manually blotted with filter paper and rapidly plunged into liquid ethane. Grids were store in liquid nitrogen, then transferred to a Gatan 626 cry-specimen holder (Gatan, Warrendale, PA) and maintained at -180 °C. Low magnification images were collected at a nominal magnification of 29,000X on a Tecnai F20 Twin transmission electron microscope (FEI, Hillsboro, OR) operating at 120kV. Digital micrographs were recorded on a Gatan US4000 CCD or Teitz XF416 camera.

**Nanoparticle Tracking Analysis (NTA):**

NTA was performed using a NanoSight NS300 (Malvern Panalytical) equipped with a high sensitivity sCMOS camera, 532 nm laser, and automatic syringe pump. NTA was used to determine the concentration and size distribution of EVs isolated from urine samples. EVs resuspended in 1x PBS were thawed on ice diluted with 1x PBS prior to injection. Camera and detection settings are provided in **Supplementary Table 1**. Videos were captured and processed using NTA 3.3 Dev Build 3.3.104 (Malvern) with 3 videos of 60 seconds per measurement. All samples were analyzed with automatic syringe movement set to 100. Concentration (EV particles/mL) and total yield (EV particles/sample) were calculated based on dilution factors and the total working solution for each isolation method.
